# Adenosine A2A Receptor (A_2A_R) agonists improve survival in K28-hACE2 mice following SARS CoV-2 infection

**DOI:** 10.1101/2022.02.25.481997

**Authors:** Barbara J. Mann, Preeti Chhabra, Mingyang Ma, Savannah G. Brovero, Marieke K. Jones, Joel Linden, Kenneth L. Brayman

## Abstract

Effective and available therapies for the treatment of COVID-19 disease are limited. Apadenoson is a highly potent selective anti-inflammatory adenosine A_2A_ receptor (A_2A_R) agonist and potential treatment option for COVID-19 patients. Apadenoson, when administered after infection with SARS CoV-2, was found to decrease weight loss, improve clinical symptoms, reduce levels of a several proinflammatory cytokines and chemokines in bronchial lavage (BAL) fluid, and promote increased survival in K18hACE2 transgenic mice. Of note, administering apadenoson after, but not prior to Covid-19 infection, caused a rapid decrease in lung viral burden. The work presented provides the foundation for further examination of these drugs as a therapy option for COVID-19.

**Summary:** Apadenoson therapy improves COVID-19 outcome

## INTRODUCTION

COVID-19, the Coronavirus disease caused by severe acute respiratory syndrome corona virus 2 (SARS CoV-2), represents a public health crisis of global proportions (Sharma et al., 2020). Declared a pandemic by the World Health Organization (WHO) on March 11, 2020, this disease continues to spread aggressively across the world. Almost all patients with COVID-19 present with lung involvement; only a subgroup of patients exhibit life-threatening complications such as Acute Respiratory Distress Syndrome (ARDS) (Xu et al., 2020). This subgroup of patients first develops severe pneumonia in the second week, accompanied by high levels of circulating cytokines (“cytokine storm”) (Li et al., 2020)(Xu et al., 2020), profound lymphopenia, eosinopenia, substantial mononuclear cell infiltration in the lungs, heart, spleen, lymph nodes and kidney (Merad and Martin, 2020), and an increased neutrophil-to-lymphocyte ratio (Merad and Martin, 2020). These symptoms eventually progress to ARDS, multi-organ failure and disseminated intravascular coagulation (DIC) (Xu et al., 2020) (Zaim et al., 2020).

Although there has been rapid development of protective vaccines, emergency-use approved therapies for COVID-19 are limited. Remdesivir, given intravenously, is a nucleoside analog that that targets the viral polymerase, was the first drug officially approved by the Federal Drug Administration for COVID-19 treatment. More recently, the FDA has also authorized “emergency use authorization” of oral Paxlovid™ for the treatment of mild to moderate COVID-19. Paxlovid™ is a combination of nirmatrelvir, which targets viral replication, and ritonavir, which extends the half-life of nirmatrelvir. Paxlovid™ is reported to reduce hospital admissions and death in high risk individuals (Mahase, 2021)(Owen et al., 2021). Administration of these antivirals is associated with shortened recovery times and less respiratory tract infection in hospitalized patients (Beigel et al., 2020). The use of remdesivir is controversial, as other studies have found that the treatment offered little benefit to hospitalized patients (WHO Solidarity Trial Consortium, 2021). However, combining remdesivir with other drugs was shown to be superior to remdesivir alone. The anti-inflammatory glucocorticosteroid dexamethosome has shown effectiveness in reducing COVID-19-related deaths, and significantly reducing hospital stays (The RECOVERY Collaborative Group, 2021), but is used at a low dose of 6 mg/day to prevent immunosuppression. Combining dexamethosome and remdesivir has been reported to reduce mortality and the need for mechanical ventilation (Benfield et al., 2021). Currently, remdesivir and dexamethasone are indicated as standard-of-care treatment for COVID-19 patients with mild-moderate disease and not requiring high flow oxygen, according to guidelines of the National Institute of Health, USA. The combination of remdesivir with baricitinib, a JAK kinase inhibitor, has proven superior to remdesivir alone by shortening hospital stays (Kalil et al., 2021). Other treatments aimed at attenuating immune pathogenesis related to cytokine storm syndromes include amongst others: IFN-γ neutralization, IL-6 receptor blockade or neutralization, B-cell ablation with rituximab, T cell–directed immunomodulation, T-cell ablation with anti-thymocyte globulincorticosteroids, IL-1 family member cytokines blockade including blockade of IL-18 binding protein, IL-1β, IL-33 receptor, and JAK inhibition (Kumar et al., 2021).

Adenosine agonists offer another potential treatment option for COVID-19 patients. Aerosolized adenosine administered twice daily was found to increase (> 30%) PaO2/FiO2-ratio in 13 out of 14 patients, despite it’s very short half live (Correale et al., 2020). In a case report two severely ill COVID-19 patients treated with inhaled adenosine exhibited a rapid improvement in lung function and one recovered (Spiess et al., 2021). Adenosine is a potent autocrine and paracrine immunosuppressive nucleoside involved in regulating both the innate and adaptive immune responses via four widely expressed G protein-coupled receptors A1, A2A, A2B and A3 (Chhabra et al., 2012). A2A or A2B adenosine receptor occupancy in most immune cells activates endogenous immunosuppressive pathways that act to reduce tissue injury and inflammation, and promotes repair by increasing oxygen supply/demand ratio, inhibiting pro-inflammatory cells and cytokines, decreasing endothelial adhesion molecule expression, and stimulating angiogenesis (Chhabra et al., 2012)(Thiel et al., 2005). Adenosine A(_2A_) receptor (A_2A_R) activation plays a critical role in providing lung protection from enhanced neutrophil accumulation, lung vascular permeability, and impairment of lung gas exchange (Chhabra et al., 2012)(Thiel et al., 2005). In asthma and COPD patients suppression of leukocyte influx into broncho-alveolar lavage (BAL) fluid and reduction of inflammatory cell activation by A_2A_R agonists contributed to immunosuppressive activity (Fozard et al., 2002). Activation of A_2A_Rs on myeloid cells attenuated cytokine release and the recruitment/ adhesion of neutrophils to pulmonary endothelial cells following LPS-induced lung injury (Reutershan et al., 2007).

Regadenoson and apadenoson (also known as ATL146e) are A_2A_R agonists. Regadenoson is used clinically for myocardial perfusion imaging because of its ability to induce coronary vasodilation without exercise and with few serious side-effects (Iskandrian et al., 2007). Apadenoson is a more potent A_2A_R selective agonist; it has been shown to improve survival in a mouse *Escherichia coli* model of sepsis and was synergistic with the antibiotic Ceftriaxone (Sullivan et al., 2004). Both regadenoson and apadenoson protect animal and human lungs from ischemia-reperfusion or transplant-injury (Ellman et al., 2008)(Lau et al., 2020). ATL146e (apadenoson) also reduced joint inflammation in combination with antibiotics in a rabbit septic arthritis model without interfering with bacterial clearance (Cohen et al., 2005). As a result of the effectiveness of apadenoson in the mouse sepsis model we tested the therapeutic efficacy of apadenoson in preventing the onset and progression of SARS CoV-2 infection in the K18-hACE2 mouse model of COVID-19. This is a model of severe COVID-19, with nearly 100% mortality by five to six days post-infection (Moreau et al., 2020). Our results indicate that apadenoson administered after SARS CoV-2 infection decreased weight loss, improved clinical symptoms, reduced several proinflammatory cytokines in the BAL fluid, and increased survival in K18hACE2 transgenic mice.

## RESULTS

### Apadenoson administration improves outcomes of SARS-CoV-2 infection in K18-hACE2 mice

K18-hACE2 mice, ranging in age from 17 to 30 weeks were divided into three groups. One group received saline (vehicle), another received apadenoson 18 hours before viral challenge (drug) and a third received apadenoson five hours post viral challenge (drug with delay). The mice were challenged intranasally with 1250 PFU SARS CoV-2. Mouse weight and clinical scores were recorded daily. Mice were euthanized when they met the criteria for euthanasia (see methods) or at the end of the experiment, which was day 12 post viral challenge. Apadenoson, given either pre- or post-challenge, improved survival and delayed the onset of symptoms (Figure 1). Apadenoson administration beginning five hours after challenge was more effective in delaying the time to death, although the overall survival was the same in both drug and drug with delay treatment groups. All mice in the vehicle treatment group met the criteria for euthanasia by day 8 (0/18 surviving); 24% (4/17) of the mice survived to the end of the experiment in the drug treatment group and 30% (5/17) survived in the drug with delay treatment group (Figure 1A). The survival curve for the drug with delay-treated group was significantly different than vehicle (p=0.0002). The survival curve for the drug-treated group did not differ significantly from the vehicle-treated group (p=0.066). The drug and drug with delay-treated groups had statistically significantly lower clinical scores and less weight loss when compared to the vehicle-treated group (Figures 1B &1C).

**Figure 1.**
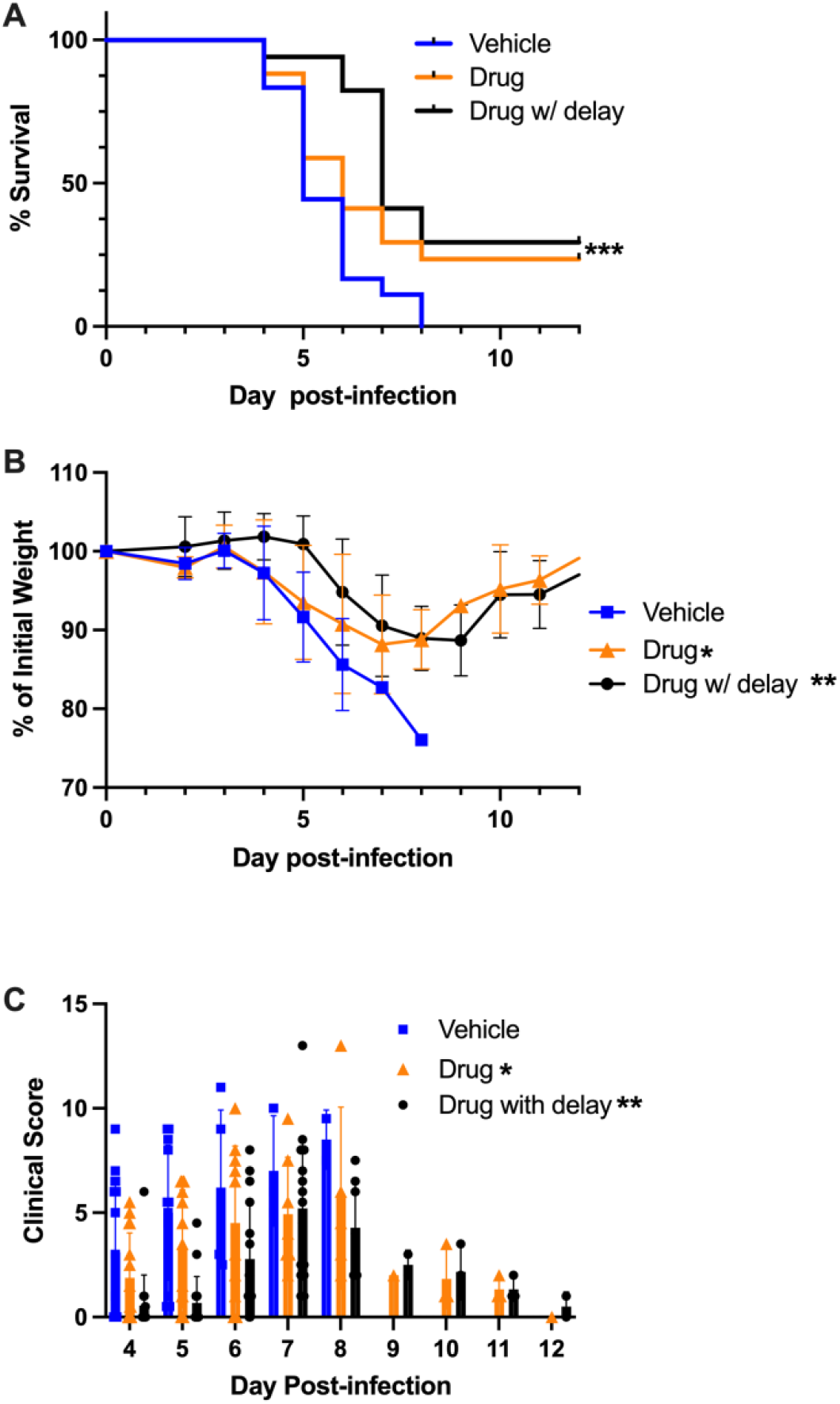
Treatment with apadenoson improved survival and delayed time to death. Mice were treated 18 hours before or five hours after infection with apadenoson delivered by an osmotic pump. Data combined from three independent experiments. A) Survival curves, used Log Rank (Mantel Cox) test, p=0.0002*** for drug with delay compared to vehicle, B) Average weight loss: P values from linear mixed model for difference in slope of vehicle compared to drug, p=0.02*, vehicle compared to drug with delay, p=0.002** and C) Clinical Scores: P values from linear mixed model for difference in slope of vehicle compared to drug, p =0.027*, vehicle compared to drug with delay, p =0.001**

### Apadenoson treatment elicits cytokine and chemokine responses that are associated with a decrease in weight loss

To assess how apadenoson treatment may be delaying the onset of severe symptoms, mice were treated as before with either vehicle (saline) or apadenoson, delivered with a five hour delay by osmotic pumps, then euthanized on day five post-infection. The weight loss and clinical scores of mice in both vehicle and drug with delay-treated animals were heterogeneous on day five and not statistically different between groups (Figure 2).

**Figure 2.**
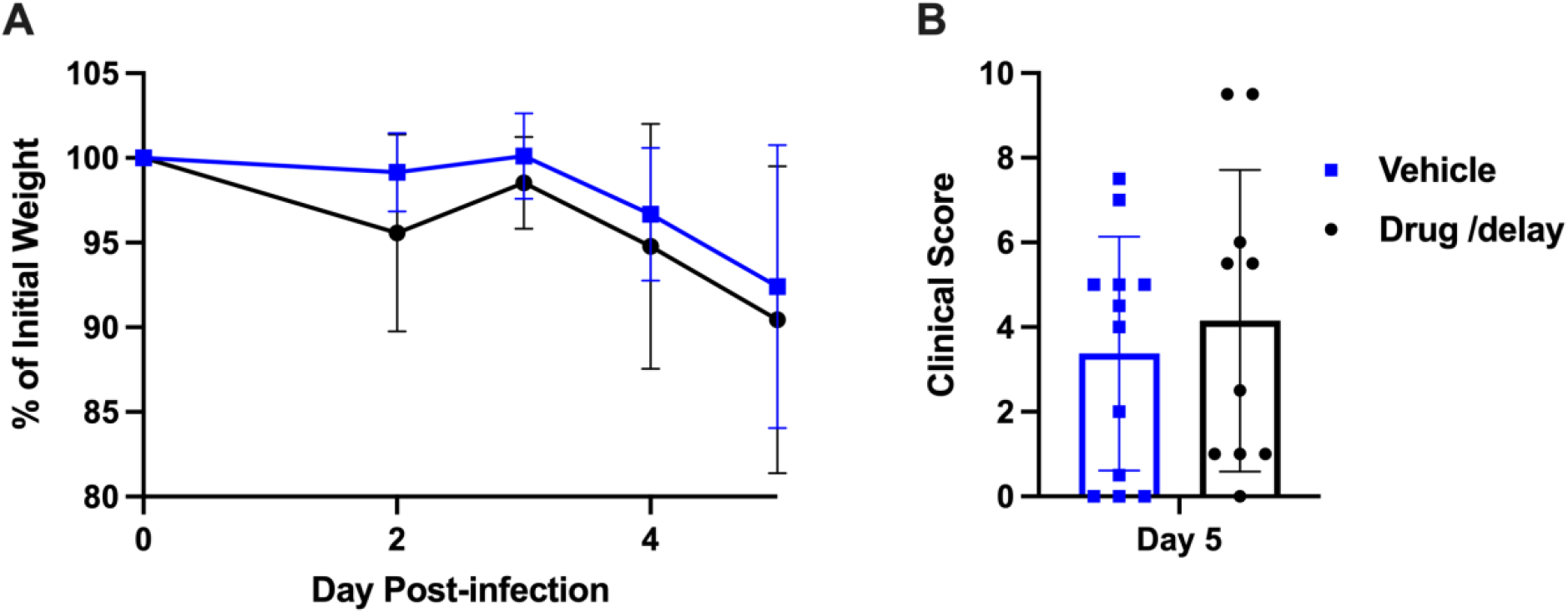
Weight loss (A) and clinical scores (B) of SARS CoV-2 infected mice treated with apadenoson (drug with delay) or saline (vehicle) were not statistically different on day five post-infection. Mean ± SD is shown. Statistically evaluated using unpaired t-test.

Cytokines and chemokines were measured in the BAL fluid using a mouse 31 plex Luminex panel. The levels of 18/31 cytokines/chemokines (IL-1a, IL-1b, IL-2, IL-4, IL-5, IL-7, IL-9, IL-10, IL-12p40, IL-12p70, IL-13, IL-15, IL-17, GM-CSF, LIX, MIP-1a, MIP2, & M-CSF) were not different between uninfected, vehicle-infected control and drug with delay-infected mice (data not shown). Uninfected mice received either saline or apadenoson via an osmotic pump; regardless of treatment these mice all had low levels of cytokines and chemokines.

The levels of 13/31 cytokines/chemokines (IP-10, MIG, MCP-1, G-CSF, Eotaxin, MIP1b, VEGF, RANTES, INF-g, IL-6, TNF-a, KC, LIF) were statistically different by ANOVA and raw p value (Figure 3). Using ANOVA p values adjusted for multiple testing, only IP-10 remained significant (p=0.022). Based on the results of Dunnett’s T3 comparisons, the levels of nine cytokines or chemokines (IP-10, MIG, MCP-1, G-CSF, Eotaxin, MIP1b, IL-6, TNF-a, KC) were higher in vehicle-infected compared to uninfected controls (p<0.05). VEGF was notable because it was lower in vehicle-treated mice compared to uninfected mice (p=0.011). Only MCP-1 and IP-10 were statistically higher in the drug with delay-infected group compared to uninfected controls (p=0.0351, and p=0.0223, respectively). The levels of eight cytokines and chemokines (MIG, G-CSF, Eotaxin, MIP1b, VEGF, INF-g, TNF-a, KC) in the drug with delay treatment group were not significantly different from uninfected mice. Taken together, these results demonstrate that treatment with apadenoson lowers the levels of a subset of proinflammatory cytokines and chemokines to baseline values.

**Figure 3.**
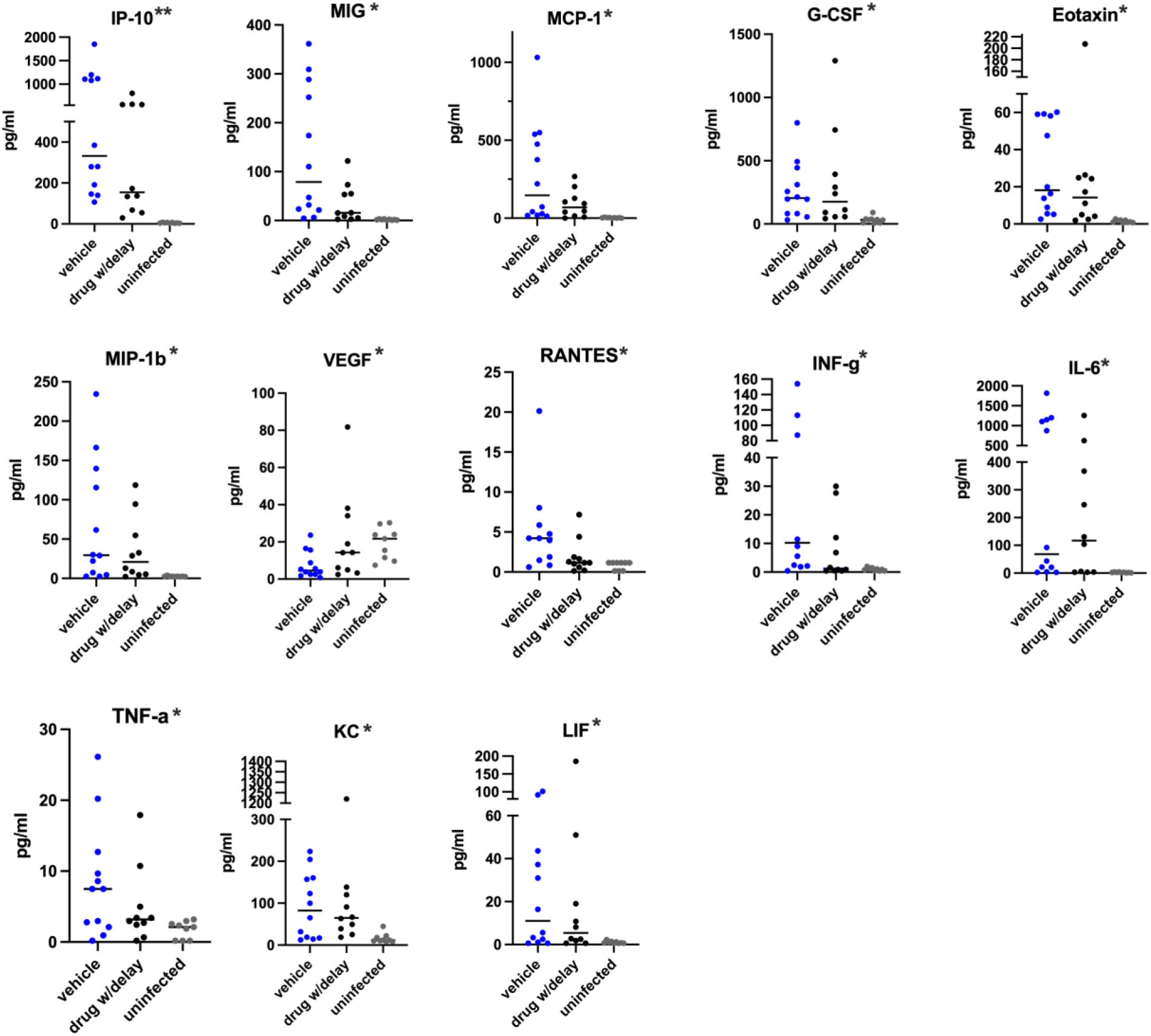
Statistically significant differences in cytokine and chemokine levels in SARS CoV-2-infected mice treated with apadenoson. Mice were euthanized on day five post-infection. Cytokine/chemokine levels were measured in BAL fluid by a Luminex. Combined results of two independent experiments. Statistical differences in cytokine/chemokine levels were analyzed using Welch’s ANOVA. Raw p values <0.05*, 0.005**

The levels of most cytokines exhibited a wide range of values within a treatment group, similar to the observed heterogeneity seen in weight loss and clinical scores. Therefore, linear relationships between cytokine levels and weight loss were examined (Figure 4). Higher levels of IP-10 and MIG were associated with less weight loss in both drug with delay- and vehicle-treated groups; p values were <0.05 in drug with delay treatment group, and <0.005 in vehicle treatment group. Lower levels of VEGF were associated with less weight loss in the drug-treatment group, p value =0.007. In the drug with delay treatment group, cytokine and chemokine levels were overall lower than in vehicle-treated mice, and with the exception of IP-10, MIG, and VEGF, not associated with the degree of weight loss. For the vehicle treatment group, higher levels of eight additional cytokines and chemokines were associated with less weight loss (Eotaxin, KC, MCP-1, MIP-1b, IL-6, INF-g, TNF-a, and LIF). There was an outlier in the drug with delay treatment group; this mouse had the most weight loss and had very high levels of five cytokines (Eotaxin, MIP-1b, IL-6, KC and LIF) that were not consistent with the associations with weight loss with mice in either group. These results suggest an extremely dysregulated immune response in this mouse, but there was no apparent explanation. In these experiments, when mice were euthanized on day five post-infection, only about half of the mice in each group had reached the established criteria for euthanasia (see methods and note clinical scores in Figure 3). With the one exception noted above, mice in either treatment group with 20% or more weight loss had low levels of cytokines. These results suggest that higher cytokine levels in the vehicle treatment group are associated with a slower progression in weight loss, and potentially a delay in time to eventual death, whereas apadenoson-treated mice exhibited a slower progression in weight loss without elevated cytokines and potentially an increased chance of survival or recovery.

**Figure. 4.**
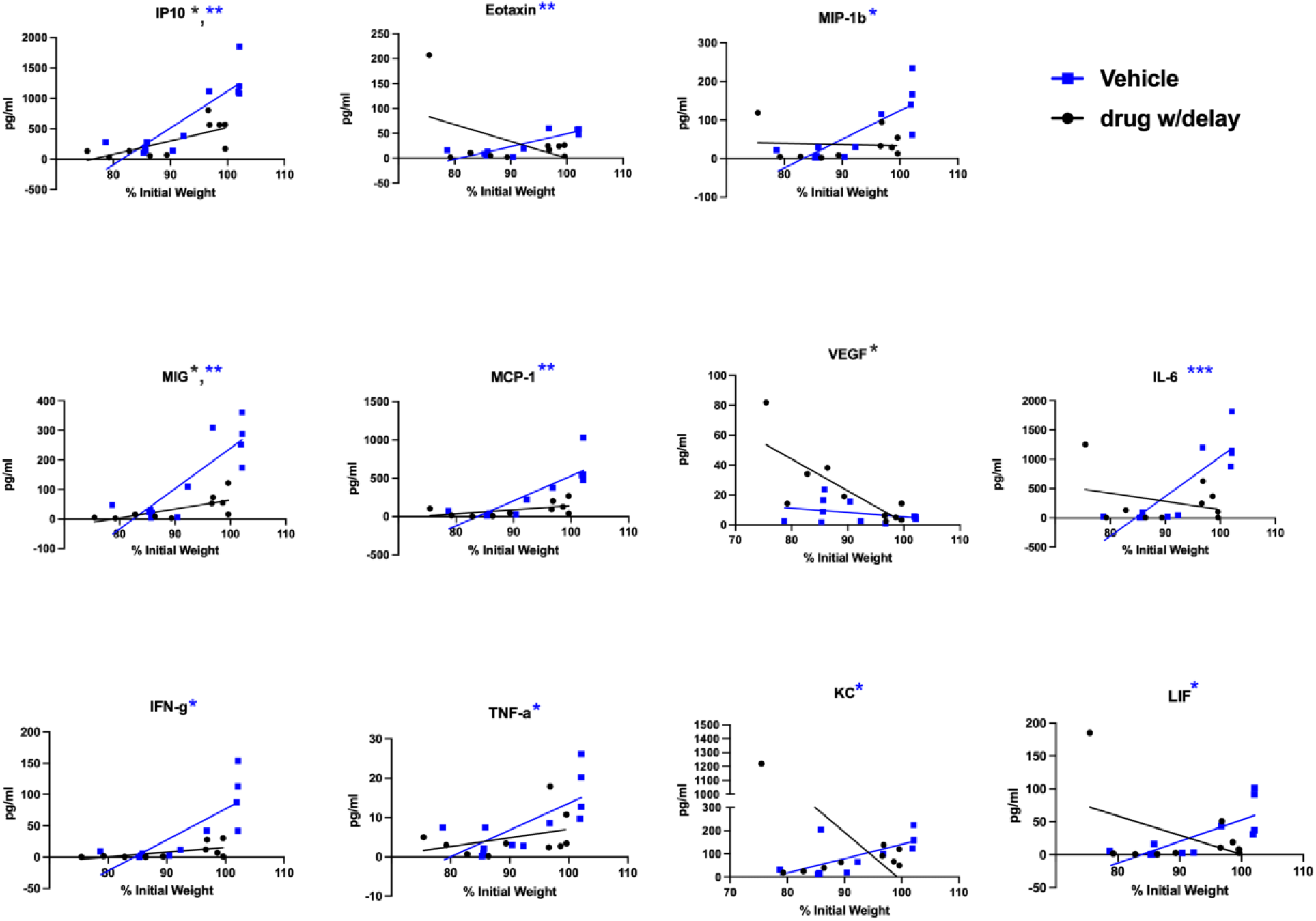
Associations of BALF cytokine and chemokine levels and weight loss. P values < 0.05 *, <0.005 ** calculated using linear regression, (blue asterisks indicate p value significance for vehicle treatment group, black asterisks for drug with delay treatment group). These results are combined from two independent experiments.

### Viral burdens and titers in the lungs in apadenoson-treated mice

To assess the effects of apadenoson treatment on viral burden, the presence of the virus in the lungs was measured by immunohistochemistry (IHC) staining with an anti-viral NP protein on day five post-infection (Figure 5A) and followed by the percentage of lung stained with the antibody. The viral burden in drug with delay-treated mice was inversely associated with weight loss, but the same was not true in the vehicle-treated group (Figure 5B, p=0.0163). Of note, in the drug with delay treatment group mice with the most weight loss (e.g., mouse #8 in Figure 5), had the least amount of viral staining in the lungs and mice with little to minimal weight loss had the highest lung burden (e.g., mouse #11, Figure 5). Viral burden and weight loss were not associated in the vehicle-treated group, supporting the characterization of COVID-19 as an inflammatory disease with dysregulated immune responses.

**Figure 5.**
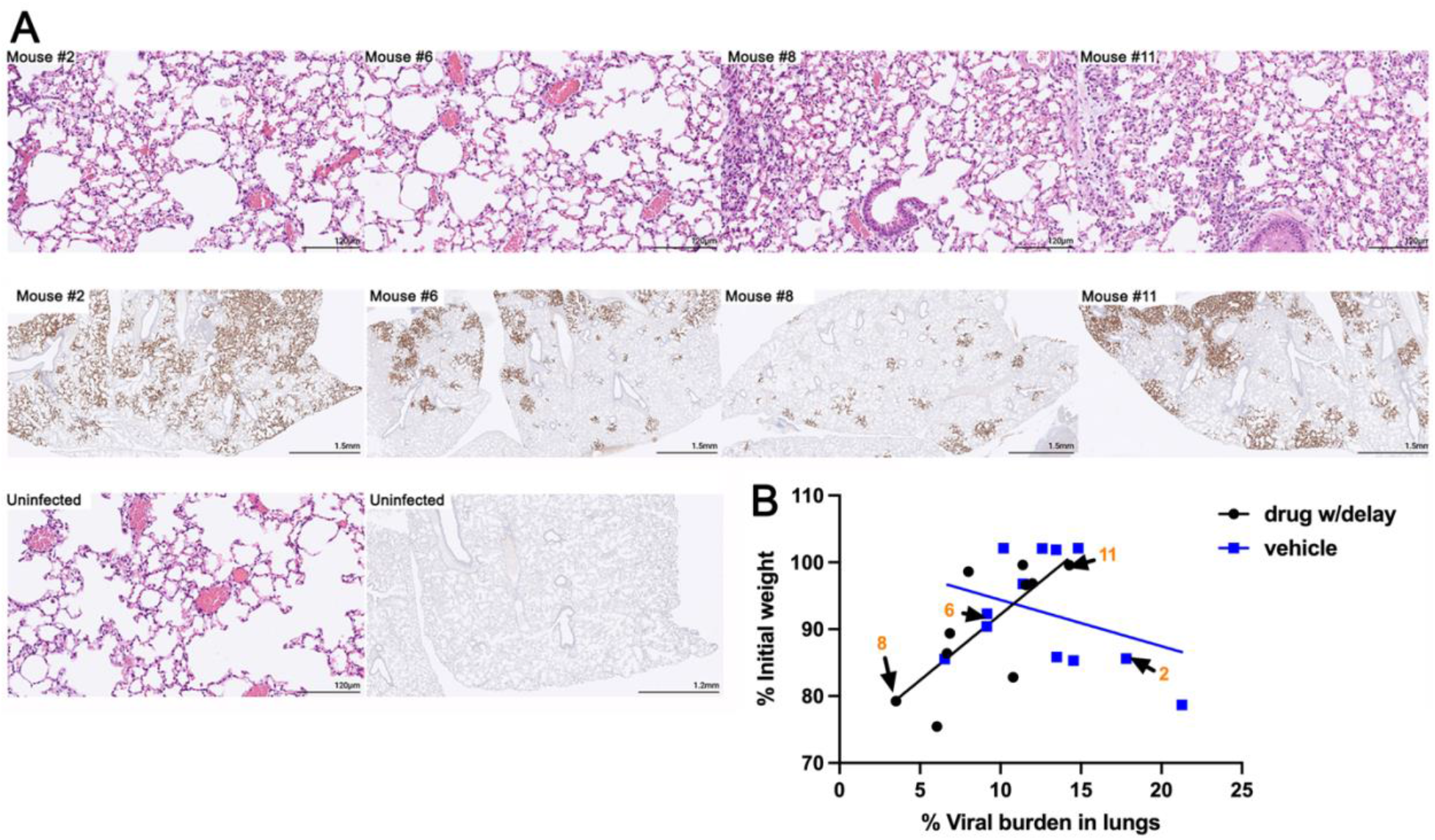
Viral burden on day five post-infection in apadenoson-treated mice negatively correlates with weight loss. A) Lungs stained with H&E or an anti-NP antibody. Mice #2 and 6 were vehicle-treated; Mice #8 and #11 were drug-treated. H&E-stained lungs are shown at a higher magnification than IHC slides. B) The weight and viral burden of mice shown in the H&E stained slides are indicated in orange numbers with black arrows on the graph. Association in drug delay group was calculated using simple linear regression, p value = 0.0163. There was no significant association in the vehicle treatment group. The percent viral burden in the lungs was determined using ImageJ.

We also assessed viral titer and burden in the lungs of mice that met the criteria for euthanasia and in mice that survived infection (after restoration of weight and no discernable clinical score, see Figure 1B & C). Weight loss and viral titer were significantly associated with apadenoson given before infection (Figure 6A, p=0.0207) based on simple logistic regression (Figure 6A), but was not significant in apadenoson given after infection (drug with delay) or in vehicle-treated mice. However, surviving mice in both drug treatment groups had viral titers below the limit of detection, whereas the vehicle-treated mice, none of which survived, all had detectable titers (Figures 6A). The average viral titer in the drug with delay treatment group trended lower, but was not significant (Figure 6B). However, the viral burden, determined by IHC staining was significantly lower in the drug with delay treatment group (Figure 6C, p=0.0012). In summary, surviving mice, which included only mice that had received apadenoson (Figure 1A), had no detectable live virus in their lungs, and lower viral burdens by IHC. There was no significant difference in the viral titers among the non-survivors regardless of treatment group. These results suggest that the pharmacological effects of apadenoson result in lowering viral burden despite down-regulating several aspects of the inflammatory response.

**Figure 6.**
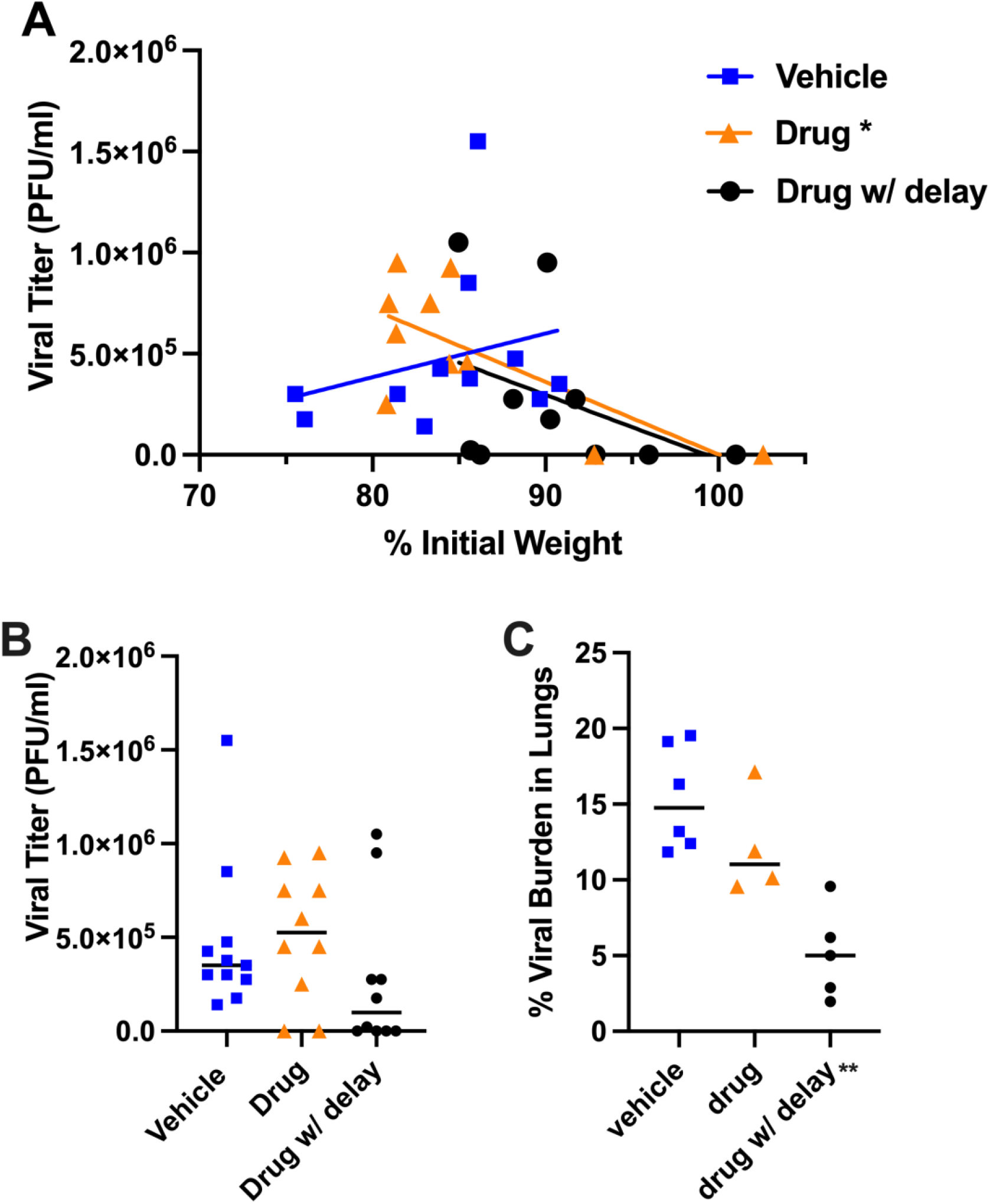
Association of viral titer and weight loss in mice and burden in lungs in apadenoson-treated mice. A) Weight loss was significantly associated with viral titer in mice treated with apadenoson before infection (drug, p=0.0207*), but not in vehicle or drug with delay treatment groups. Slopes were calculated using simple linear regression. Virus was not detected in surviving mice (See Figure 1). B) Viral titers of lung homogenates trended lower in drug with delay-treated mice but were not significant using Welch’s ANOVA. C) Viral burdens, as measured by IHC, were significantly lower in drug with delay-treated mice (p=0.0012**) using Welch’s ANOVA.

## Discussion

Immune system activation is essential for killing COVID-19 and other viruses, but prolonged hyperinflammation and immune dysregulation contribute to poor outcomes in COVID-19 disease (Merad and Martin, 2020)(Coperchini et al., 2020). Adenosine agonists represent attractive therapies for COVID-19, because they limit several elements of immune responses and promote tissue repair (Chhabra et al., 2012). We used the transgenic K18hACE2 mouse, which expresses the human ACE2 receptor, as a model of severe COVID-19 disease. This model shares several characteristics of severe COVID-19 disease including impaired lung function, high levels of proinflammatory cytokines and chemokines, and immune cell infiltration of the lungs (Yinda et al., 2021)(Winkler et al., 2020). We have shown in this mouse model that apadenoson, administered five hours after infection, is capable of delaying the progression of symptoms and death.

Mice treated after infection with apadenoson had dampened immune responses compared to vehicle controls, as measured by reduced levels of a subset of cytokines and chemokines that were similar to baseline levels in uninfected mice. Nine cytokines and chemokines were increased in vehicle-treated mice (IP-10, MIG, MCP-1, G-CSF, Eotaxin, MIP1b INF-g, TNF-a, KC) compared to uninfected controls (Figure 3). These cytokine and chemokine profiles in the vehicle-treated infected mice are similar to other reports in K18hACE2 mice (Yinda et al., 2021)(Winkler et al., 2020) and confirm the inflammatory nature of COVID-19 infection in this model.

To explore the impact of apadenoson on cytokine/chemokine levels further these levels were correlated with percent weight loss. On day five, when half of the mice in each group met the criteria for euthanasia (Figure 2), there was an association between low levels of cytokines and increased weight loss in vehicle-treated mice (Figure 4). In the drug with delay treatment group only higher levels of IP10, and MIG were associated with less weight loss (Figure 4). The remaining cytokines in the drug with delay treatment group were low, regardless of the amount of weight loss. The sole exception was VEGF, which was higher in animals with increased weight loss, though this association was only significant in drug with delay-treated mice. VEGF promotes vascular permeability, which can lead to tissue damage, and is elevated in COVID-19 patients (Huang et al., 2020). A promising preliminary COVID-19 treatment trial with bevacizumab, an anti-VEGF monoclonal antibody, suggests that this treatment may be beneficial. (Pang et al., 2021). Although we did not do a time course of cytokine induction, our results suggest that in this model cytokine levels peak before severe weight loss and symptomatic disease occur, but then drop as the disease progresses.

Apadenoson treatment also influenced viral burden in lung. In mice that were euthanized on day five post-infection, which in this model is the day when weight loss and adverse clinical symptoms become apparent, there was an inverse relation between viral lung burden and weight loss in the drug with delay-treated mice, but no strong association in the vehicle-treated mice (Figure 5). Of note, mice in both groups that had not yet exhibited weight loss on day five post-infection tended to have viral lung burdens between 10-15%. Based on the drug treatment trials in Figure 1, we would expect that all vehicle-treated mice would have succumbed to infection, while some of the apadenoson-treated mice would have survived. Therefore, we next examined the viral titers and burden in the lungs of mice that were euthanized or recovered (Figure 6). While viral titers were not strongly related to weight loss, the surviving apadenoson-treated mice did not have detectable titers, and viral burden was significantly lower with mice given apadenoson after infection, suggesting that apadenoson can also lead to clearance of the virus. Overall, these findings suggest that the use of anti-inflammatory drugs such as apadenoson to limit inflammation during cytokine storm is beneficial and remarkably, and may even reduce viral burden in the lung. These data support the use and further exploration of apadenoson, or other adenosine agonists, as supportive care for COVID-19 patients.

## Methods

### Infection and treatment

Twenty-four hours prior to infecting Tg(K18-*hACE2*)2Prlmn (Jackson Laboratories) male mice with SARS CoV-2, primed 7-day Alzet® osmotic pumps (Durect, Cuperton, CA) containing saline or drug were implanted subcutaneously (McCray et al., 2007). In the drug with delay treatment group drug delivery was delayed by 24 hours by adding 2.5 cm long tubing to the pump. Apadenoson was delivered at a rate of 1.5 μg/kg/hr. Uninfected littermates received saline or drug with delay via osmotic pumps. Mice were challenged with 1250 plaque forming units (PFU) of Hong Kong/VM20001061/2020 (NR-52282, BEI Resources) via intranasal route under ketamine/xylazine anesthesia. Mice were examined twice daily for clinical symptoms and scored using the following criteria: weight loss (0-5, in 5% of initial weight loss increments), reduced activity (0-3), ruffled fur appearance and hunched posture (0-2), and an eye closure (0-2). Mice were euthanized when weight was <80% of initial weight or when they scored a maximum in two symptom categories. This experiment was repeated three times, with six animals per treatment group each time. One mouse in the drug and one in the drug with delay did not survive anesthesia. Statistical differences in weight loss and clinical scores by treatment group were determined using linear mixed models accounting for day post-infection and the day the mouse died (Ime4 and ImerTest packages (Bates et al., 2015)(Kuznetsova et al., 2017) in R (R Core Team 2021). For mice surviving to the end of the experiment (day 12 post-infection), day of death =15 was used. All mouse work was approved by the University’s Institutional Animal Care and Use Committee and all procedures were performed in the University’s certified animal Biosafety Level Three laboratory, which is fully accredited by the Association for the Assessment and Accreditation of Laboratory Animal Care, International (AAALAC).

### BAL Cytokines

BAL fluid was collected with 700 μL of PBS and cells were removed via centrifugation. The BAL fluid was analyzed by a Luminex Magpix by the UVA Flow Cytometry Core facility using a Mouse 31 plex panel. Statistical changes in cytokine levels were analyzed using Welch’s ANOVA followed by Dunnett’s T3 multiple comparisons (Graphpad Prism 9.0). P values were adjusted for multiple testing using the Holm-Sidak method.

### Viral Lung titers

Titers were determined as described in Moreau *et al*.(Moreau et al., 2020). Briefly, lungs were homogenized in serum-free DMEM. Titers were determined by infecting Vero C1008, Clone E6 (ATCC CRL-1586) with serial dilutions of the homogenate. After a two hour incubation, the diluted homogenates were replaced with a liquid overlay of DMEM, 2.5% FBS containing 1.2% Avicel PH-101 (Sigma Aldrich, St. Louis, MO) and incubated at 37°C, 5% CO_2_. After three days, the overlay was removed, wells were fixed with 10% formaldehyde, and stained with 0.1% crystal violet to visualize plaques. Plaques were counted and PFUs were calculated according to the following equation: Average # of plaques/dilution factor × volume of diluted virus added to the well.

### Histology

Tissues were fixed in 10% formalin. To visualize virus in the lungs, slides were stained with SARS-CoV2 specific anti-nucleoprotein antibody (Cat. No. 9099, ProSci, Poway, CA) as per manufacturer’s instructions and then scanned at 20X magnification. The percentage of lung tissue infected with virus was calculated (ImageJ, version 1.53K).

## Acknowledgements

This work was supported by a grant from the Manning Family Foundation. B.J. Mann, S.G. Brovero, and M. Jones have no conflict of interest. P. Chhabra, J. Linden, and K.L. Brayman have a patent pending on the use of A_2A_R agonists for the treatment of COVIID-19. We would like to acknowledge and thank the University of Virginia’s Research Histology Core and Biorepository and Tissue Research Facility for preparation and staining of histology slides.

## References

Bates, D., Mächler, M., Bolker, B., Walker, S., 2015. Fitting Linear Mixed-Effects Models Using lme4. J. Stat. Softw. 67. https://doi.org/10.18637/jss.v067.i01

Beigel, J.H., Tomashek, K.M., Dodd, L.E., Mehta, A.K., Zingman, B.S., Kalil, A.C., Hohmann, E., Chu, H.Y., Luetkemeyer, A., Kline, S., Lopez de Castilla, D., Finberg, R.W., Dierberg, K., Tapson, V., Hsieh, L., Patterson, T.F., Paredes, R., Sweeney, D.A., Short, W.R., Touloumi, G., Lye, D.C., Ohmagari, N., Oh, M., Ruiz-Palacios, G.M., Benfield, T., Fätkenheuer, G., Kortepeter, M.G., Atmar, R.L., Creech, C.B., Lundgren, J., Babiker, A.G., Pett, S., Neaton, J.D., Burgess, T.H., Bonnett, T., Green, M., Makowski, M., Osinusi, A., Nayak, S., Lane, H.C., 2020. Remdesivir for the Treatment of Covid-19 — Final Report. N. Engl. J. Med. 383, 1813–1826. https://doi.org/10.1056/NEJMoa2007764

Benfield, T., Bodilsen, J., Brieghel, C., Harboe, Z.B., Helleberg, M., Holm, C., Israelsen, S.B., Jensen, J., Jensen, T.Ø., Johansen, I.S., Johnsen, S., Lindegaard, B., Lundgren, J., Meyer, C.N., Mohey, R., Pedersen, L.M., Nielsen, H., Nielsen, S.L., Obel, N., Omland, L.H., Podlekareva, D., Poulsen, B.K., Ravn, P., Sandholdt, H., Starling, J., Storgaard, M., Søborg, C., Søgaard, O.S., Tranborg, T., Wiese, L., Christensen, H.R., 2021. Improved Survival Among Hospitalized Patients With Coronavirus Disease 2019 (COVID-19) Treated With Remdesivir and Dexamethasone. A Nationwide Population-Based Cohort Study. Clin. Infect. Dis. ciab536. https://doi.org/10.1093/cid/ciab536

Chhabra, P., Linden, J., Lobo, P., Douglas Okusa, M., Lewis Brayman, K., 2012. The Immunosuppressive Role of Adenosine A2A Receptors in Ischemia Reperfusion Injury and Islet Transplantation. Curr. Diabetes Rev. 8, 419–433. https://doi.org/10.2174/157339912803529878

Cohen, S.B., Leo, B.M., Baer, G.S., Turner, M.A., Beck, G., Diduch, D.R., 2005. An adenosine A2A receptor agonist reduces interleukin-8 expression and glycosaminoglycan loss following septic arthrosis. J. Orthop. Res. 23, 1172–1178. https://doi.org/10.1016/j.orthres.2005.01.015

Coperchini, F., Chiovato, L., Croce, L., Magri, F., Rotondi, M., 2020. The cytokine storm in COVID-19: An overview of the involvement of the chemokine/chemokine-receptor system. Cytokine Growth Factor Rev. 53, 25–32. https://doi.org/10.1016/j.cytogfr.2020.05.003

Correale, P., Caracciolo, M., Bilotta, F., Conte, M., Cuzzola, M., Falcone, C., Mangano, C., Falzea, A.C., Iuliano, E., Morabito, A., Foti, G., Armentano, A., Caraglia, M., De Lorenzo, A., Sitkovsky, M., Macheda, S., 2020. Therapeutic effects of adenosine in high flow 21% oxygen aereosol in patients with Covid19-pneumonia. PLOS ONE 15, e0239692. https://doi.org/10.1371/journal.pone.0239692

Ellman, P.I., Reece, T.B., Law, M.G., Gazoni, L.M., Singh, R., Laubach, V.E., Linden, J., Tribble, C.G., Kron, I.L., 2008. Adenosine A2A Activation Attenuates Nontransplantation Lung Reperfusion Injury. J. Surg. Res. 149, 3–8. https://doi.org/10.1016/j.jss.2007.08.008

Fozard, J.R., Ellis, K.M., Villela Dantas, M.F., Tigani, B., Mazzoni, L., 2002. Effects of CGS 21680, a selective adenosine A2A receptor agonist, on allergic airways inflammation in the rat. Eur. J. Pharmacol. 438, 183–188. https://doi.org/10.1016/S0014-2999(02)01305-5

Huang, C., Wang, Y., Li, X., Ren, L., Zhao, J., Hu, Y., Zhang, L., Fan, G., Xu, J., Gu, X., Cheng, Z., Yu, T., Xia, J., Wei, Y., Wu, W., Xie, X., Yin, W., Li, H., Liu, M., Xiao, Y., Gao, H., Guo, L., Xie, J., Wang, G., Jiang, R., Gao, Z., Jin, Q., Wang, J., Cao, B., 2020. Clinical features of patients infected with 2019 novel coronavirus in Wuhan, China. The Lancet 395, 497–506. https://doi.org/10.1016/S0140-6736(20)30183-5

Iskandrian, A., Bateman, T., Belardinelli, L., Blackburn, B., Cerqueira, M., Hendel, R., Lieu, H., Mahmarian, J., Olmsted, A., Underwood, S., 2007. Adenosine versus regadenoson comparative evaluation in myocardial perfusion imaging: Results of the ADVANCE phase 3 multicenter international trial. J. Nucl. Cardiol. 14, 645–658. https://doi.org/10.1016/j.nuclcard.2007.06.114

Kalil, A.C., Patterson, T.F., Mehta, A.K., Tomashek, K.M., Wolfe, C.R., Ghazaryan, V., Marconi, V.C., Ruiz-Palacios, G.M., Hsieh, L., Kline, S., Tapson, V., Iovine, N.M., Jain, M.K., Sweeney, D.A., El Sahly, H.M., Branche, A.R., Regalado Pineda, J., Lye, D.C., Sandkovsky, U., Luetkemeyer, A.F., Cohen, S.H., Finberg, R.W., Jackson, P.E.H., Taiwo, B., Paules, C.I., Arguinchona, H., Erdmann, N., Ahuja, N., Frank, M., Oh, M., Kim, E.-S., Tan, S.Y., Mularski, R.A., Nielsen, H., Ponce, P.O., Taylor, B.S., Larson, L., Rouphael, N.G., Saklawi, Y., Cantos, V.D., Ko, E.R., Engemann, J.J., Amin, A.N., Watanabe, M., Billings, J., Elie, M.-C., Davey, R.T., Burgess, T.H., Ferreira, J., Green, M., Makowski, M., Cardoso, A., de Bono, S., Bonnett, T., Proschan, M., Deye, G.A., Dempsey, W., Nayak, S.U., Dodd, L.E., Beigel, J.H., 2021. Baricitinib plus Remdesivir for Hospitalized Adults with Covid-19. N. Engl. J. Med. 384, 795–807. https://doi.org/10.1056/NEJMoa2031994

Kumar, S., Çalışkan, D.M., Janowski, J., Faist, A., Conrad, B.C.G., Lange, J., Ludwig, S., Brunotte, L., 2021. Beyond Vaccines: Clinical Status of Prospective COVID-19 Therapeutics. Front. Immunol. 12, 752227. https://doi.org/10.3389/fimmu.2021.752227

Kuznetsova, A., Brockhoff, P.B., Christensen, R.H.B., 2017. lmerTest Package: Tests in Linear Mixed Effects Models. J. Stat. Softw. 82. https://doi.org/10.18637/jss.v082.i13

Lau, C.L., Beller, J.P., Boys, J.A., Zhao, Y., Phillips, J., Cosner, M., Conaway, M.R., Petroni, G., Charles, E.J., Mehaffey, J.H., Mannem, H.C., Kron, I.L., Krupnick, A.S., Linden, J., 2020. Adenosine A2A receptor agonist (regadenoson) in human lung transplantation. J. Heart Lung Transplant. 39, 563–570. https://doi.org/10.1016/j.healun.2020.02.003

Li, X., Geng, M., Peng, Y., Meng, L., Lu, S., 2020. Molecular immune pathogenesis and diagnosis of COVID-19. J. Pharm. Anal. 10, 102–108. https://doi.org/10.1016/j.jpha.2020.03.001

Mahase, E., 2021. Covid-19: Pfizer’s paxlovid is 89% effective in patients at risk of serious illness, company reports. BMJ n2713. https://doi.org/10.1136/bmj.n2713

McCray, P.B., Pewe, L., Wohlford-Lenane, C., Hickey, M., Manzel, L., Shi, L., Netland, J., Jia, H.P., Halabi, C., Sigmund, C.D., Meyerholz, D.K., Kirby, P., Look, D.C., Perlman, S., 2007. Lethal Infection of K18-hACE2 Mice Infected with Severe Acute Respiratory Syndrome Coronavirus. J. Virol. 81, 813–821. https://doi.org/10.1128/jvi.02012-06

Merad, M., Martin, J.C., 2020. Pathological inflammation in patients with COVID-19: a key role for monocytes and macrophages. Nat. Rev. Immunol. 20, 355–362. https://doi.org/10.1038/s41577-020-0331-4

Moreau, G.B., Burgess, S.L., Sturek, J.M., Donlan, A.N., Petri, W.A., Mann, B.J., 2020. Evaluation of K18-hACE2 Mice as a Model of SARS-CoV-2 Infection. Am. J. Trop. Med. Hyg. 103, 1215–1219. https://doi.org/10.4269/ajtmh.20-0762

Owen, D.R., Allerton, C.M.N., Anderson, A.S., Aschenbrenner, L., Avery, M., Berritt, S., Boras, B., Cardin, R.D., Carlo, A., Coffman, K.J., Dantonio, A., Di, L., Eng, H., Ferre, R., Gajiwala, K.S., Gibson, S.A., Greasley, S.E., Hurst, B.L., Kadar, E.P., Kalgutkar, A.S., Lee, J.C., Lee, J., Liu, W., Mason, S.W., Noell, S., Novak, J.J., Obach, R.S., Ogilvie, K., Patel, N.C., Pettersson, M., Rai, D.K., Reese, M.R., Sammons, M.F., Sathish, J.G., Singh, R.S.P., Steppan, C.M., Stewart, A.E., Tuttle, J.B., Updyke, L., Verhoest, P.R., Wei, L., Yang, Q., Zhu, Y., 2021. An oral SARS-CoV-2 M ^pro^ inhibitor clinical candidate for the treatment of COVID-19. Science 374, 1586–1593. https://doi.org/10.1126/science.abl4784

Pang, J., Xu, F., Aondio, G., Li, Y., Fumagalli, A., Lu, M., Valmadre, G., Wei, J., Bian, Y., Canesi, M., Damiani, G., Zhang, Yuan, Yu, D., Chen, J., Ji, X., Sui, W., Wang, B., Wu, S., Kovacs, A., Revera, M., Wang, H., Jing, X., Zhang, Ying, Chen, Y., Cao, Y., 2021. Efficacy and tolerability of bevacizumab in patients with severe Covid-19. Nat. Commun. 12, 814. https://doi.org/10.1038/s41467-021-21085-8

R Core Team, n.d. R: A language and environment for statistical computing. R Foundation for Statistical Computing, Vienna, Austria.

Reutershan, J., Cagnina, R.E., Chang, D., Linden, J., Ley, K., 2007. Therapeutic Anti-Inflammatory Effects of Myeloid Cell Adenosine Receptor A2a Stimulation in Lipopolysaccharide-Induced Lung Injury. J. Immunol. 179, 1254–1263. https://doi.org/10.4049/jimmunol.179.2.1254

Sharma, A., Tiwari, S., Deb, M.K., Marty, J.L., 2020. Severe acute respiratory syndrome coronavirus-2 (SARS-CoV-2): a global pandemic and treatment strategies. Int. J. Antimicrob. Agents 56, 106054. https://doi.org/10.1016/j.ijantimicag.2020.106054

Spiess, B.D., Sitkovsky, M., Correale, P., Gravenstein, N., Garvan, C., Morey, T.E., Fahy, B.G., Hendeles, L., Pliura, T.J., Martin, T.D., Wu, V., Astrom, C., Nelson, D.S., 2021. Case Report: Can Inhaled Adenosine Attenuate COVID-19? Front. Pharmacol. 12, 676577. https://doi.org/10.3389/fphar.2021.676577

Sullivan, G.W., Fang, G., Linden, J., Scheld, W.M., 2004. A _2A_ Adenosine Receptor Activation Improves Survival in Mouse Models of Endotoxemia and Sepsis. J. Infect. Dis. 189, 1897–1904. https://doi.org/10.1086/386311

The RECOVERY Collaborative Group, 2021. Dexamethasone in Hospitalized Patients with Covid-19. N. Engl. J. Med. 384, 693–704. https://doi.org/10.1056/NEJMoa2021436

Thiel, M., Chouker, A., Ohta, A., Jackson, E., Caldwell, C., Smith, P., Lukashev, D., Bittmann, I., Sitkovsky, M.V., 2005. Oxygenation Inhibits the Physiological Tissue-Protecting Mechanism and Thereby Exacerbates Acute Inflammatory Lung Injury. PLoS Biol. 3, e174. https://doi.org/10.1371/journal.pbio.0030174

WHO Solidarity Trial Consortium, 2021. Repurposed Antiviral Drugs for Covid-19 — Interim WHO Solidarity Trial Results. N. Engl. J. Med. 384, 497–511. https://doi.org/10.1056/NEJMoa2023184

Winkler, E.S., Bailey, A.L., Kafai, N.M., Nair, S., McCune, B.T., Yu, J., Fox, J.M., Chen, R.E., Earnest, J.T., Keeler, S.P., Ritter, J.H., Kang, L.-I., Dort, S., Robichaud, A., Head, R., Holtzman, M.J., Diamond, M.S., 2020. SARS-CoV-2 infection of human ACE2-transgenic mice causes severe lung inflammation and impaired function. Nat. Immunol. 21, 1327–1335. https://doi.org/10.1038/s41590-020-0778-2

Xu, Z., Shi, L., Wang, Y., Zhang, J., Huang, L., Zhang, C., Liu, S., Zhao, P., Liu, H., Zhu, L., Tai, Y., Bai, C., Gao, T., Song, J., Xia, P., Dong, J., Zhao, J., Wang, F.-S., 2020. Pathological findings of COVID-19 associated with acute respiratory distress syndrome. Lancet Respir. Med. 8, 420–422. https://doi.org/10.1016/S2213-2600(20)30076-X

Yinda, C.K., Port, J.R., Bushmaker, T., Offei Owusu, I., Purushotham, J.N., Avanzato, V.A., Fischer, R.J., Schulz, J.E., Holbrook, M.G., Hebner, M.J., Rosenke, R., Thomas, T., Marzi, A., Best, S.M., de Wit, E., Shaia, C., van Doremalen, N., Munster, V.J., 2021. K18-hACE2 mice develop respiratory disease resembling severe COVID-19. PLOS Pathog. 17, e1009195. https://doi.org/10.1371/journal.ppat.1009195

Zaim, S., Chong, J.H., Sankaranarayanan, V., Harky, A., 2020. COVID-19 and Multiorgan Response. Curr. Probl. Cardiol. 45, 100618. https://doi.org/10.1016/j.cpcardiol.2020.100618

